# Mycobiome analysis of electronic cigarettes reveals a reservoir of pathogenic yeasts

**DOI:** 10.1101/2025.08.01.668078

**Authors:** Katy Deitz, Zifan Zhao, Yana Goddard, Arantxa V. Lazarte, Borna Mehrad, Jason Smith

## Abstract

Research on the health impacts of e-cigarettes has focused on non-infectious manifestations. Given their enclosed plastic design and temperature fluctuations, we hypothesized that e-cigarettes are colonized by pathogenic microbes, thereby contributing to lung disease in users. Using sequencing and culture techniques of the devices and mouthwash of 25 users, we found only a small subset of mouthpieces to contain bacteria, whereas most were abundantly colonized with fungi that were distinct from the oral mycobiota, including the genera *Rhodotorula, Aureobasidium, Cystobasidium*, and *Meyerozyma*. Chronic exposure to the most frequently isolated pathogen, *Cystobasidium minutum*, resulted in mucus hypersecretion and obstructive lung disease in mice, characteristics of chronic bronchitis. We conclude that e-cigarettes are frequently colonized with fungal organisms capable of causing lung disease.

## Introduction

E-cigarette use has increased exponentially since their introduction in 2007, with recent estimates at 8.1 million adults and more than 2.5 million youth users in the United States[1–3]. Compared to tobacco smoking, our understanding of the health effects of e-cigarette use is in its infancy.

E-cigarette use has been linked to a form of acute lung injury, e-cigarette or vaping product use-associated lung injury (EVALI), and several chronic respiratory illnesses, including asthma, bronchitis, and bronchiolitis, but the underlying biological mechanisms of these illnesses are only beginning to be defined[2,4]. Previous research has explored the toxicity induced by chemical constituents of the vehicle of e-cigarettes, and their effect on oral and periodontal bacterial microbiota[5–10]. In contrast, the microbial communities colonizing e-cigarette devices themselves, and their potential health consequences, have not been studied systematically.

In the healthcare setting, plastic devices can be colonized by thermotolerant yeasts capable of causing nosocomial infections[11]. We therefore hypothesized that e-cigarettes are colonized by pathogenic microbes, thereby contributing to lung disease in users.

## Results

### Sample characteristics

We enrolled 25 e-cigarette users, most of whom used e-cigarettes daily, had disposable devices, and did not clean their devices. About a third of the subjects reported respiratory symptoms (Table 1).

**Table 1.**
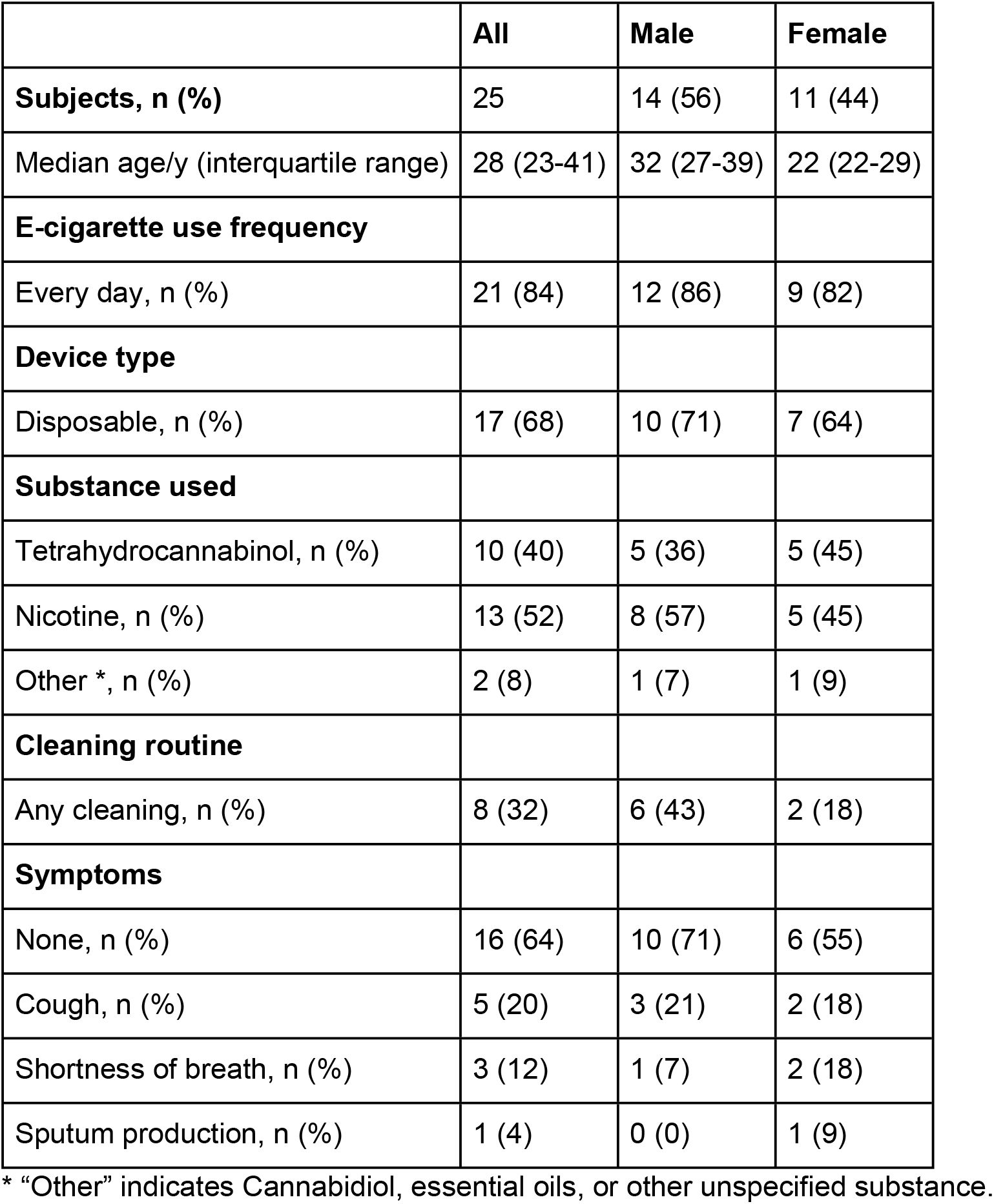
Summary Demographics of study subjects

|  | All | Male | Female |
| --- | --- | --- | --- |
| **Subjects, n (%)** | 25 | 14 (56) | 11 (44) |
| Median age/y (interquartile range) | 28 (23-41) | 32 (27-39) | 22 (22-29) |
| **E-cigarette use frequency** |  |  |  |
| Every day, n (%) | 21 (84) | 12 (86) | 9 (82) |
| **Device type** |  |  |  |
| Disposable, n (%) | 17 (68) | 10 (71) | 7 (64) |
| **Substance used** |  |  |  |
| Tetrahydrocannabinol, n (%) | 10 (40) | 5 (36) | 5 (45) |
| Nicotine, n (%) | 13 (52) | 8 (57) | 5 (45) |
| Other *, n (%) | 2 (8) | 1 (7) | 1 (9) |
| **Cleaning routine** |  |  |  |
| Any cleaning, n (%) | 8 (32) | 6 (43) | 2 (18) |
| **Symptoms** |  |  |  |
| None, n (%) | 16 (64) | 10 (71) | 6 (55) |
| Cough, n (%) | 5 (20) | 3 (21) | 2 (18) |
| Shortness of breath, n (%) | 3 (12) | 1 (7) | 2 (18) |
| Sputum production, n (%) | 1 (4) | 0 (0) | 1 (9) |

\* "Other" indicates Cannabidiol, essential oils, or other unspecified substance.

### Culture

Among the e-cigarette samples, 52% produced fungal colonies in culture, resulting in 35 isolates which were successfully amplified using ITS rDNA (Table 2). Potential human pathogens represented 80% of the isolates and were identified as: *Rhodotorula taiwanensis, Rhodotorula glutinis, Cystobasidium minutum, Mucor circinelloides, Papiliotrema flavescens, Aureobasidum melanogenum, Aureobasidium pullulans*, and *Meyerozyma guilliermondii* (Figure 1B) [11–25]. Of these, *C. minutum* had the highest occurrence and was selected for *in vivo* studies.

**Table 2.** Fungal isolates identified in this study with Sanger sequencing.

| Sample # | Best BLASTn Match | Media | % Identity | E-value |
| --- | --- | --- | --- | --- |
| VS-1 | *R. taiwanensis* (LC498459.1) | CHROMagar | 100.00% | 0 |
|  | *R. taiwanensis* (LC498459.1) | Yeast Agar | 100.00% | 0 |
|  | *R. glutinis* (EF194846.1) | MEA+ | 99.52% | 0 |
| VS-2 | *R. taiwanensis* (LC498459.1) | MEA+ | 100.00% | 0 |
|  | *R. taiwanensis* (LC498459.1) | Yeast | 100.00% | 0 |
|  | *R. taiwanensis* (NR157462.1) | APDA | 99.64% | 0 |
| VS-3 | *R. taiwanensis* (LC498459.1) | MEA+ | 100.00% | 0 |
|  | *R. taiwanensis* (LC498459.1) | Yeast Agar | 100.00% | 0 |
| VS-4 | *C. minutum* (LC529381.1) | APDA | 99.82% | 0 |
|  | *C. minutum* (LC497828.1) | CHROMagar | 100.00% | 0 |
| VS-5 | *C. minutum* (LC497828.1) | APDA | 100.00% | 0 |
|  | *C. minutum* (LC497828.1) | MEA+ | 100.00% | 0 |
|  | *C. minutum* (MT256942.1) | MEA | 99.60% | 0 |
| VS-6 | *C. minutum* (KY107435.1) | APDA | 100.00% | 0 |
|  | *C. minutum* (KY107435.1) | CHROMagar | 100.00% | 0 |
| VS-7 | *C. minutum* (KY107435.1) | APDA | 100.00% | 0 |
|  | *C. minutum* (KY107435.1) | CHROMagar | 100.00% | 0 |
| VS-8 | *C. minutum* (OM238182.1) | CHROMagar | 99.46% | 0 |
|  | *C. minutum* (KY107435.1) | MEA+ | 100.00% | 0 |
|  | *C. minutum* (KY107435.1) | APDA | 100.00% | 0 |
|  | *C. minutum* (KY107435.1) | MEA | 100.00% | 0 |
| VS-11 | *C. minutum* (KY107435.1) | MEA+ | 100.00% | 0 |
|  | *C. minutum* (MT254989.1) | APDA | 100.00% | 0 |
|  | *C. minutum* (MT254989.1) | CHROMagar | 100.00% | 0 |
|  | *C. minutum* (KY107435.1) | Yeast agar | 100.00% | 0 |
| VS-14 | *M. circinelloides*<br>(MT603954.1) | CHROMagar | 99.78% | 0 |
|  | *C. minutum* (AF190010) | MEA+ | 99.47% | 0 |
| VS-15 | *P. flavescens* (MG815868.1) | CHROMagar | 100.00% | 0 |
|  | *P. flavescens* (MT447858.1) | Yeast Agar | 98.95% | 0 |
| VS-19 | *A. melanogenum*<br>(MT623572.1) | APDA | 100.00% | 0 |
|  | *A. melanogenum*<br>(MT444123.1) | CHROMagar | 99.64% | 0 |
|  | *A. pullulans* (MT237304.1) | MEA+ | 100.00% | 0 |
| VS-24 | *M. guilliermondii*<br>(MT247857.1) | CHROMagar | 100.00% | 0 |
|  | *M. guilliermondii*<br>(MT247857.1) | MEA+ | 100.00% | 0 |
|  | *M. guilliermondii*<br>(MT247857.1) | Yeast Agar | 100.00% | 0 |

**Figure 1.**
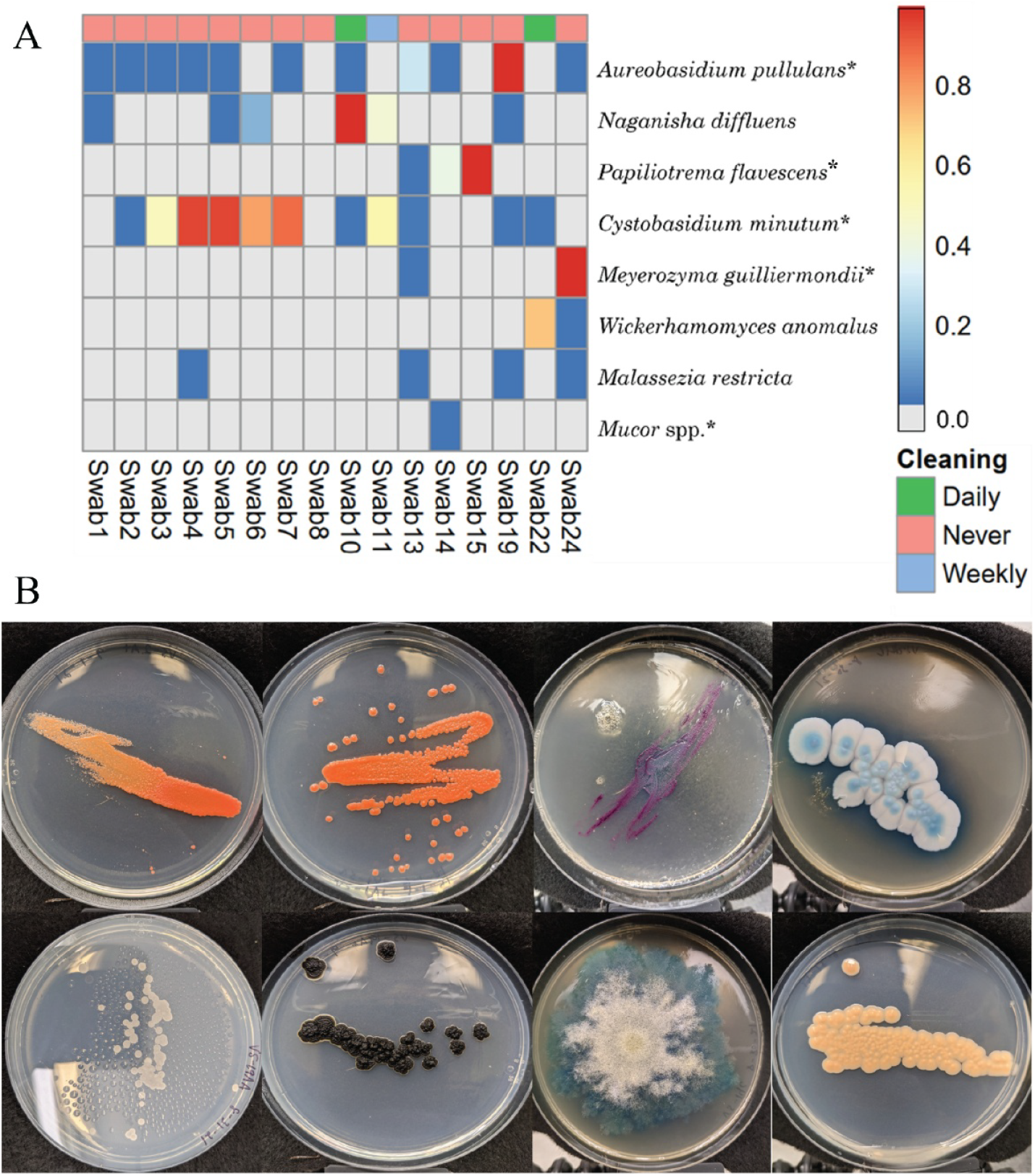
(A) Heatmap showing the relative abundance of cultured fungi and all the potential pathogens from the ASV data. The fungi detected from the NGS result that were also cultured indicated with an asterisk. (B) Representative fungal cultures recovered from e-cigarette mouthpieces in this study at 18 days. All colonies were grown at 37 ° C in the dark. From left to right, top row: *Rhodotorula taiwanensis*; *Rhodotorula glutinis*; *Papilotrema flavescens*; *Meyerozyma guilliermondii*. Second row: *Aureobasidium melanogenum*; *Aureobasidium pullulans*; *Mucor circinelloides*; *Cystobasidum minutum*

### Animal studies

After daily intrapulmonary challenge with *C. minutum* for 14 days, mice appeared well and did not lose weight. On baseline pulmonary function testing, mice challenged with *C. minutum* displayed normal inspiratory capacity, but increased breathing compared to animals challenged with saline vehicle (Figure 2A-B). After methacholine administration, mice challenged with *C. minutum* displayed both increased total respiratory resistance and elastance, indicating airway hyperresponsiveness (Figure 2C-D).

**Figure 2.**
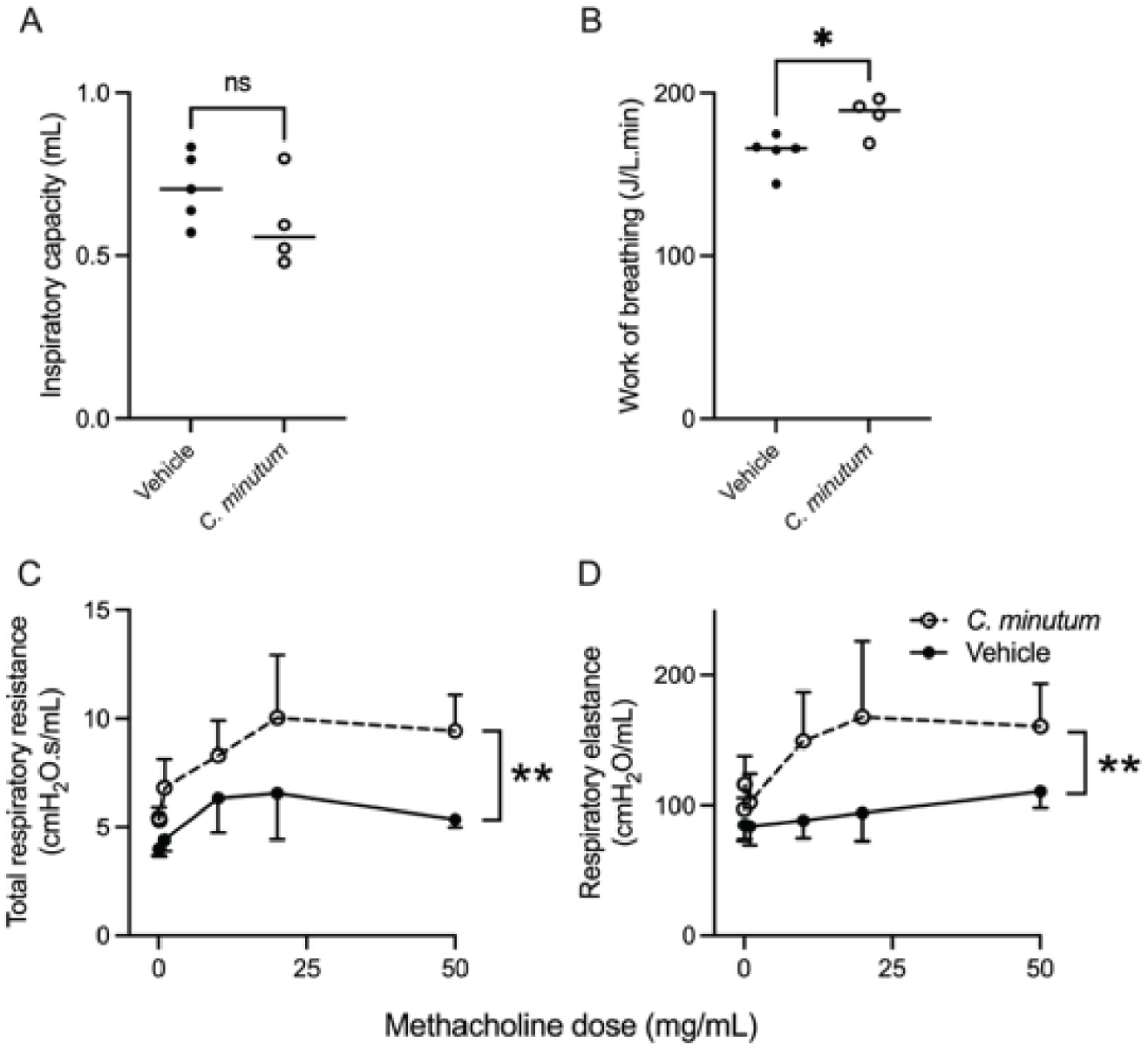
Lung physiology in animals after intra-pulmonary challenge with *C. minutum* or PBS vehicle. Dots in panels A-B represent individual animals, and lines represent medians. Panels C-D show mean and SEM of 4-5 animals per group after challenge with different doses of methacholine. ns, no significant difference; ^*^, p <0.05; ^**^, p<0.01.

Histologically, animal lungs challenged with the vehicle appeared normal (Figure 3A, C-E). In contrast, animal lungs challenged with *C. minutum* demonstrated marked mucus accumulation in the airway lumen, airway remodeling, and goblet cell hyperplasia, associated with remnants of yeast cells associated with epithelial cells (Figure 3B, F-H). MUC5AC protein level was increased in animal lungs challenged with *C. minutum* (Figure 4).

**Figure 3.**
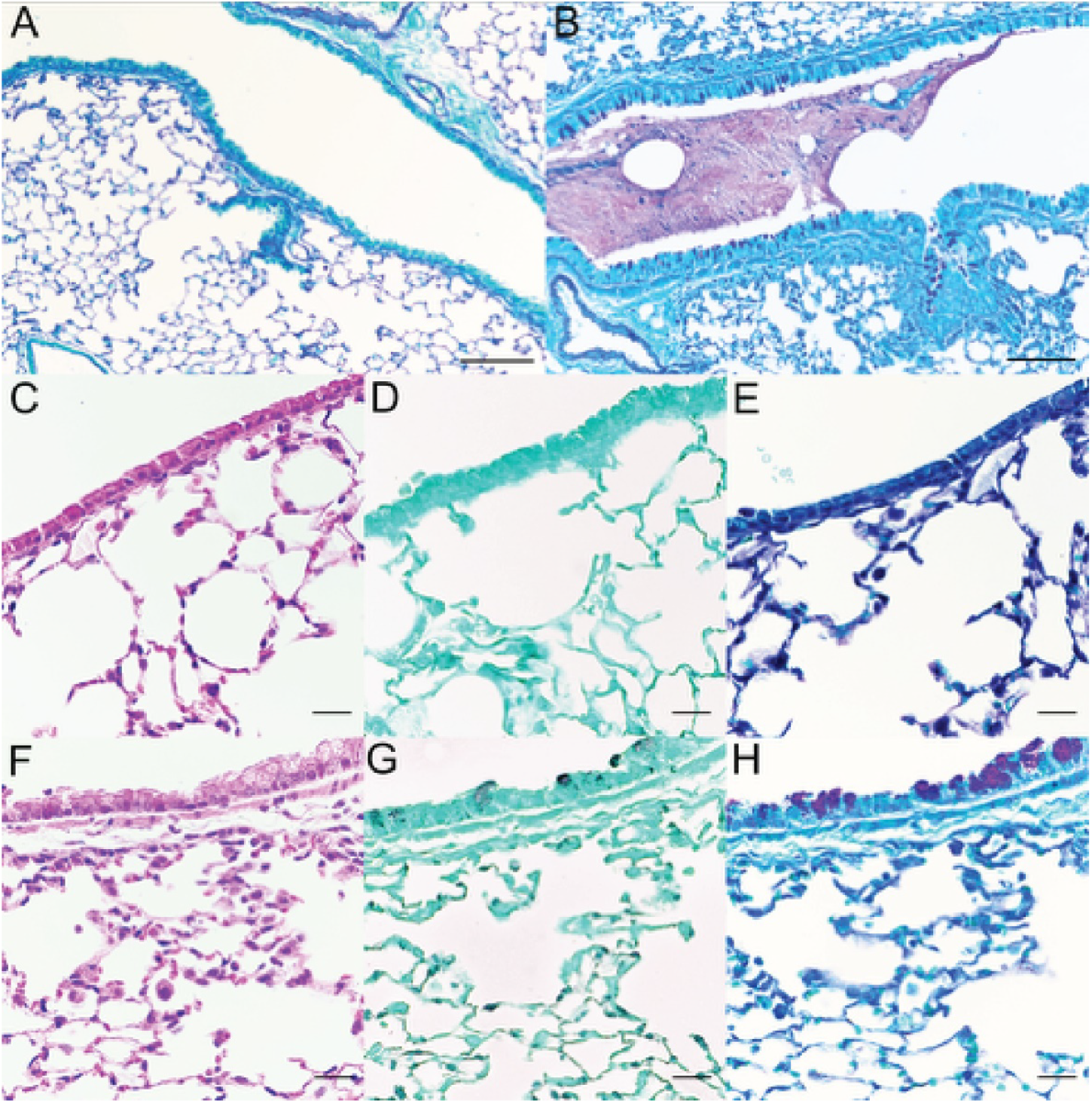
Lung histology of wildtype mice, challenged daily with intrapulmonary vehicle control (panel A, C-E) or *Cystobasidium* (panels B, F-H) for 2 weeks. As compared to control animals, animals challenged with *Cystobasidium* showed airway mucus and goblet cell hyperplasia on periodic acid-Schiff stain (panels A, B, E, H), a thickened airway epithelial basement membrane and mild inflammation on hematoxylin and eosin stain (panels C and F); and remnants of yeast cells associated with epithelial cells on Gomori methenamine silver stain (panels D and G). Scale bars are 100µm in panels A-B (original magnification 100x) and 20µm in panels C-H (original magnification 400x).

**Figure 4.**
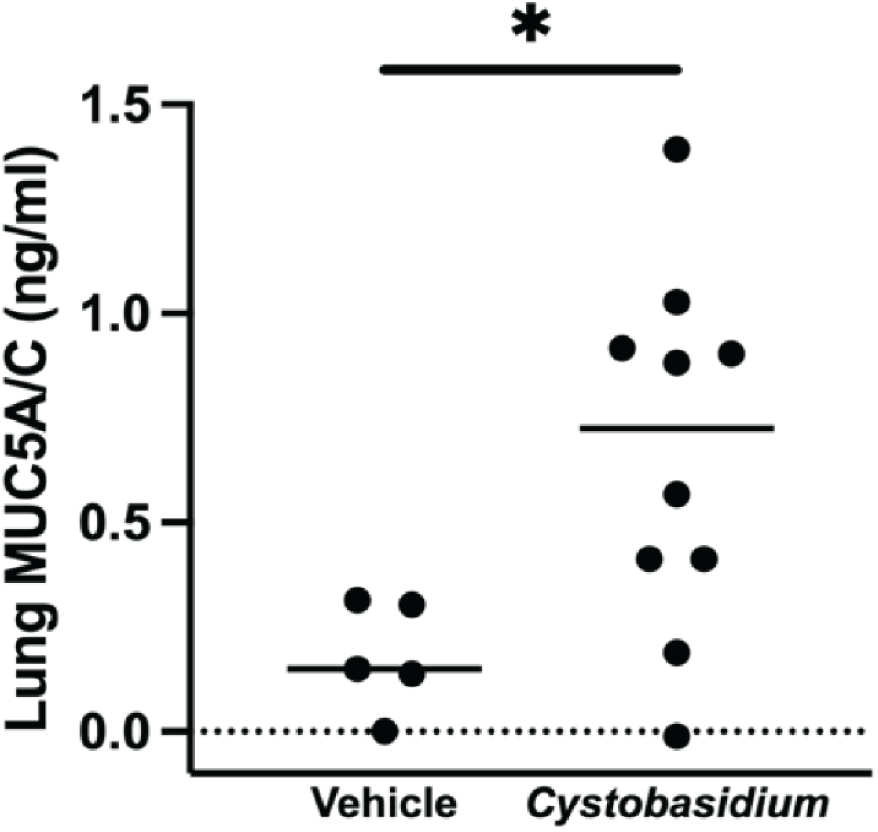
Lung MUC5AC protein concentration in animals after intra-pulmonary challenge with *C. minutum* or PBS vehicle. Dots represent individual animals, and lines represent medians. ^*^, p <0.05.

Finally, we sought to recover *C. minutum* from lungs challenged with the organism. Among 10 animals challenged with *C. minutum*, the lungs of only one animal yielded fungal growth. The morphology was characteristic of *C. minutum*, and its DNA was positively identified against GenBank archived sequences, showing a 99.5% match to the ITS region and 100% match with the D1/D2 region.

## Discussion

E-cigarettes are battery-powered atomizer devices that deliver an aerosol of nicotine or other drugs, flavoring chemicals, solvent carriers, and myriad impurities to the user’s lungs[26]. Given their variable device characteristics and the heterogeneous composition of the e-liquid, health effects remain poorly understood. The best described health effects of these devices are acute physiological effects measured after controlled experimental exposure in the laboratory, and severe and distinctive sporadic clinical syndromes, such as EVALI [27]. Evidence about their chronic health effects is limited. Most of the published literature in this field is phenomenological, defining the epidemiologic association between use and respiratory symptoms, defining acute physiologic effects, and establishing a causal link to cancers in experimental animals[28–30]. A smaller literature has focused on potential mechanistic contribution of e-cigarettes to disease, for example linking exposure to the e-liquid solvent carrier to disruption of surfactant homeostasis and predisposition to acute lung injury[31]. A subset of the latter has sought to assess the potential health effects of e-cigarettes by altering the user’s microbiota, indicating that exposure results in oral dysbiosis[9,32–34].

Little is known about the microbial content of the devices themselves. Our study adds to this literature by documenting the extensive colonization of device mouthpieces by diverse microbial communities that are distinct from the users’ oral microbiota. We specifically found most devices to be colonized by fungi, the origin of which are unlikely to have come from the user’s mouth. Surveys of the oral mycobiome have shown *Candida* to be the most reported with the highest abundance in terms of biomass; however, studies show a wide variability between oral mycobiome populations among non-*Candida* species isolated in lower abundance[35–37].

The fungal species identified in our study inhabit a wide range of niches from agricultural, environmental, and possibly skin[37–40]. The environment of the mouthpiece may select for fungal taxa with certain traits, such as thermotolerance and ability to colonize plastics. Most of the fungal species we isolated have previously been reported as capable of causing human disease, and several species, including *C. minutum, R. glutinis*, and *M. circinelloides* exhibit thermotolerance, a critical trait for environmental fungi to be considered risks to human health[11,13,14,16,18,19,21,23,25,29]. Similarly, several of the isolates (*M. guilliermondii, C. minutum, P. flavescens, R. glutinis*) resist or have variable responses to antifungal medications[16]. We speculate that some traits that enable colonization and adaptability to the microenvironment of the e-cigarette device are shared among pathogenic taxa.

*C. minutum* is a basidiomycetous yeast, previously belonging to the *R. minuta* clade[41]. Fungi in the genus *Cystobasidium* (Lagerheim) Neuhoff (1924) are dimorphic, presenting as mycoparasites on coprophilous Ascomycetes and as yeasts in the environment when the host is absent[40]. Yeast cells typically produce carotenoids, ranging in color from coral to orange. The production of mycosporine-glutaminol-glucoside, an ultraviolet radiation absorbing compound, as well as its ability to exhibit antibacterial activity may also contribute to *C. minutum*’s survival in unfavorable conditions[42,43].

Repeated pulmonary exposure of immunocompetent animals to *C. minutum* resulted in obstructive lung physiology together with some features of asthma and COPD, the most common human obstructive lung diseases. Animals challenged with *C. minutum* developed prominent features of mucus hypersecretion, namely goblet cell hyperplasia, increased lung mucin protein, and the development of airway mucus plugs. In contrast, the airways exhibited only modest inflammation. Mucus hypersecretion is a canonical feature of asthma and COPD, where it correlates with disease severity, and may represent a target for therapy[44–48]. Mucus hypersecretion has been reported as a feature of e-cigarette use, and is a feature of lung epithelial response to inhaled fungi[49–52]. Recent evidence indicates e-cigarette use correlates with COPD development independent of cigarette smoking[53,54]. In this context, our data suggests the hypothesis that fungal contamination of e-cigarette devices contributes to disease development.

In conclusion, this study documents the microbial communities colonizing in e-cigarette mouthpieces to consist predominantly of fungal species distinct from the user’s oral microbiota. *C. minutum*, is a thermotolerant yeast capable of biofilm formation and antifungal resistance[40,55]. When introduced into the lungs, this organism did not colonize the respiratory epithelium but caused prominent mucus hypersecretion and obstructive lung disease. Our study suggests several avenues for future research. First, our study requires replication by other groups and in other locations. Second, the relationship between the colonizing yeasts and other components of e-liquid aerosol in mediating lung disease in experimental animals may be important. Finally, our data suggests that culture-independent methods of assessing the microbial content of the respiratory tract of e-cigarette users, and correlation to respiratory symptoms, are of interest.

## Materials and Methods

### Study design and subject population

Study design and survey methods were reviewed and approved by the University of Florida Internal Review Board (IRB202000440). Sterile environmental sampling swabs were used (EnviroMax, Puritan, Guilford, Maine, USA) to obtain samples from e-cigarette mouthpieces. Each participant rinsed their mouth with 20 ml of Crest Scope mouthwash (Procter & Gamble, Cincinnati, Ohio, USA) for 30 seconds, which was collected into a sterile collection vial and kept on ice until transferred to storage at 4 °C.

### Fungal cultures of e-cigarette devices

Ten mL of sterile double-distilled water was added to each swab collection vial, vortexed for 30 seconds, and 20 μl were plated and spread with a sterile glass spreader on various media (See Supplementary Methods): Plates were incubated at 37°C for 3 weeks.

### Sanger sequencing and identification of isolates

DNA extractions were completed using Extract-N-Amp (Sigma Aldrich, St. Louis, Missouri, USA) minikit solution and protocol. DNA amplification utilized primers ITS1F and ITS4 targeting the ITS region of the rDNA, as well as primers NL1 and NL4 targeting the D1/D2 region of the large (28S) ribosomal subunit[56,57]. Amplicons were purified using ExoSAP-IT (Applied Biosystems, Waltham, Massachusetts, USA) and were sequenced at GeneWiz (Azenta, Burlington, Massachusetts, USA). Sequence reads were aligned and contigs edited and trimmed using Geneious (Biomatters Ltd., Auckland, New Zealand) to generate a consensus sequence. Using the BLASTn tool, final consensus sequences were compared against references in the GenBank database. GenBank accession numbers are available in Supplementary Table 1.

### Preparation of C. minutum suspension for in vivo delivery

An isolate of *C. minutum* (VS-8) was cultured in 40 ml of yeast medium broth (10 g yeast extract, 5 g NaCl, 10 g of peptone, 1 L DDI H20) for 48 hours at 37°C in a MaxQ4000 orbital shaker (Thermo Fisher). The yeast forms were pelleted at 1800 rpm for 10 minutes, washed 3 times with PBS, and enumerated under a hemocytometer.

### In vivo studies and tissue harvest

Animal studies were performed under institutionally approved protocols. Male and female 8-week-old C57BL/6J mice were purchased from Jackson Labs (Bar Harbor, Maine, USA) and maintained in specific-pathogen-free conditions in the animal care facility at the University of Florida. Forty μl of sterile PBS containing 10^6^ *C. minutum* yeast forms or sterile PBS alone was intranasally delivered to the animals daily for 14 days. Mice were monitored daily for clinical status and weight. On day 15, animals were anesthetized with tribromoethanol and subjected to lung physiology measurements using a FlexiVent system (Scireq, Montreal, Quebec, Canada), as previously described[58].

### Measurements on harvested lung tissues

Samples were processed for histology as previously described[59,60]. ELISA was performed using a commercial kit for MUC5AC (Novus Biologicals, Centennial, CO, USA) following the manufacturers’ instructions.

### Statistical analysis

Data were analyzed in Prism (version 9, GraphPad Software, San Diego, California, USA). For all statistical tests, *p* values <0.05 were considered statistically significant.

## Data and materials availability

All data are available in the manuscript or the supplementary materials.

